# FASSO: An AlphaFold based method to assign functional annotations by combining sequence and structure orthology

**DOI:** 10.1101/2022.11.10.516002

**Authors:** Carson M Andorf, Shatabdi Sen, Rita K Hayford, John L Portwood, Ethalinda K Cannon, Lisa C Harper, Jack M Gardiner, Taner Z Sen, Margaret R Woodhouse

## Abstract

Methods to predict orthology play an important role in bioinformatics for phylogenetic analysis by identifying orthologs within or across any level of biological classification. Sequence-based reciprocal best hit approaches are commonly used in functional annotation since orthologous genes are expected to share functions. The process is limited as it relies solely on sequence data and does not consider structural information and its role in function. Previously, determining protein structure was highly time-consuming, inaccurate, and limited to the size of the protein, all of which resulted in a structural biology bottleneck. With the release of AlphaFold, there are now over 200 million predicted protein structures, including full proteomes for dozens of key organisms. The reciprocal best structural hit approach uses protein structure alignments to identify structural orthologs. We propose combining both sequence- and structure-based reciprocal best hit approaches to obtain a more accurate and complete set of orthologs across diverse species, called Functional Annotations using Sequence and Structure Orthology (FASSO). Using FASSO, we annotated orthologs between five plant species (maize, sorghum, rice, soybean, Arabidopsis) and three distance outgroups (human, budding yeast, and fission yeast). We inferred over 270,000 functional annotations across the eight proteomes including annotations for over 5,600 uncharacterized proteins. FASSO provides confidence labels on ortholog predictions and flags potential misannotations in existing proteomes. We further demonstrate the utility of the approach by exploring the annotation of the maize proteome.

## Introduction

Ortholog detection in comparative genomics and bioinformatics is of fundamental importance in many fields of biology, particularly in the annotation of newly sequenced organisms, functional genomics, gene organization in species, evolutionary studies of biological systems, and phylogenomic analyses [1]. Central features of core cellular processes such as cell cycle progression, DNA replication, and transcription are often conserved among phylogenetic groups as diverse as yeast, humans, and plants. This extensive conservation has driven the use of model organisms to study conserved processes that are more difficult to assay in complex systems.

Sequence similarity, in particular, has emerged as a critical tool in predicting functional properties. In fact, most annotations of newly sequenced genomes, including gene prediction, gene function, and regulatory elements, are based on sequence similarity with other genomes that have annotated functions [2–4]. Perhaps the most common approach for inferring putative orthology is sequence-based Reciprocal Best Hits (RBH), [5–7]. Essentially, if protein *a* in genome *A* finds protein *b* as its best, highest-scoring match in sequence similarity in genome *B*; and protein *b* finds protein *a* as its best match in sequence similarity in genome *A*, they are RBH and thus identified to be “orthologs.” It is important to note that this sequence-based approach only points to high-sequence similarity, and because it lacks an evolutionary analysis, it only provides possible orthologous pairs. For practical purposes, however, we will call these pairs orthologous. The most commonly used program for finding RBH sequence matches is blastp [8]. Additional programs such as blat [9], lastal [10], Diamond [11], and MMseqs2 [12]. Each method has trade-offs in performance, with Diamond offering a good compromise in speed, sensitivity, and quality [13, 14].

Along with sequence similarity, structural similarity among proteins is used to infer similar functions, although the relationship can be complex [15, 16]. A protein’s tertiary structure largely determines its functional properties, and its structure depends on its amino acid sequence. Therefore, protein sequence similarity is used as a predictor of functional similarity, though it is less accurate than structural similarity because different amino acid sequences can yield similar structures, and similar sequences can sometimes produce different structures [17, 18]. Because of these drawbacks, there is a limit to using sequence as a proxy for structure. As established in several studies [19, 20], structural similarity in pairs with a sequence identity of 20–35%, often referred to as the “twilight zone”, is considerably less common; only fewer than ten percent of protein pairs with sequence identity below 25% have similar structures. At the same time, the “twilight zone” is characterized by an explosion of false negatives [20], which means that many dissimilar sequences appear to be structurally homologous. Consequently, protein structure and function do not always appear related to sequence, causing sequence similarity-based predictions to identify more false negatives than expected [15]. Thus, sequence similarity is a less reliable predictor of protein function than structural similarity [18, 21].

Nevertheless, sequence alignment, based on the matching of residue identity, is widely used in similarity studies due to the fact that establishing sequence similarity between sequences is an easier problem to solve and fast string-comparison algorithms were already developed in the field of computer science. Therefore the jump from strings to sequence did not take much effort. As a result, a wealth of methodologies have been developed for the alignment and comparison of sequences, in contrast to structures. In addition, it is easier and cheaper to experimentally determine protein sequences than structures, as substantiated by, a negligibly small number of solved protein structures (∼44,000 entries in the Protein Database (PDB) [22]) compared to the number of known protein sequences (over 200 million protein sequences in UniProt database [23]). However, improved computational resources and modern tools for structural alignment are expanding opportunities to efficiently implement structural comparisons across proteins [24].

Recent breakthroughs [25, 26] via AlphaFold [27] and other technologies including RoseTTAFold, OmegaFold, and ESMFold [28–31] can now predict accurate protein structures for any amino acid sequence. These predictions dramatically increase the scope of comparative structural studies [32]. Protein structures are now available for over 230 million UniProt proteins at the AlphaFold Protein Structure Database [33], including proteomes with few experimentally determined structures. For example, the crop species maize, sorghum, rice, and soybean have 235, 24, 329, and 169 experimentally-determined protein structures, respectively, in PDB (October 2022), yet AlphaFold has now computationally predicted protein structures across these species’ whole proteomes. Most plant species are also sparsely annotated with experimentally validated functions and, therefore, rely on inference for functional annotations. Developing methods to make better predictions, find novel orthologs, and provide confidence levels for ortholog predictions would greatly benefit the plant research community.

With the avalanche of predicted protein structures, new and improved tools are emerging [32, 34–39] to quickly and reliably align protein structures and assign quality and expectation scores. FoldSeek [35] and FATCAT [34] are two examples. FoldSeek uses a novel structural alphabet based on tertiary interactions. FoldSeek is up to 40,000 times faster than other approaches [35]. FoldSeek finds alignments between a protein query and target database within minutes and can compare all-vs-all alignments between proteomes within a day on a high-performance computing (HPC) environment. FATCAT aligns the backbone atoms (C^α^) and allows for flexible alignments by allowing twists around pivot points in the structures. The trade-off is that FATCAT is more resource intensive and not always practical for all-vs-all proteome comparisons. A reciprocal best structure hit approach (RBSH) using FoldSeek detected potential novel orthologs between human, worm, fruit fly, and yeast [21]. Here in this paper, we extend this approach by aggregating predictions from a sequence-based approach (Diamond) with two structure-based approaches (FoldSeek and FATCAT) to identify structural orthologs, assign functional annotations, and flag conflicting predictions. Our method, named Functional Annotations using Sequence and Structure Orthology (FASSO), identifies structural orthologs and transfers functional annotations for eight diverse proteomes that include five plants (maize, sorghum, rice, soybean, and Arabidopsis) and three well-annotated outgroups (human, *S. cerevisiae*, and *S. pombe*).

With the continued advancement of sequencing technologies, it is now possible to sequence any genome of interest, identify genes, and predict protein structures. FASSO leverages both protein sequence and structure information to improve the accuracy of proteome-scale functional annotation across a wide range of taxa by increasing the number of high-quality predictions and removing potential erroneous annotations. The ability to quickly and accurately compare and annotate proteomes is paramount for functional gene research, planning experiments, making important research decisions, and generating the formation of new hypotheses.

## Materials and Methods

### FASSO: Functional Annotations using Sequence and Structure Orthology

FASSO combines the results of three methods, Diamond, FoldSeek, and FATCAT, to identify structural orthologs and infer functional annotations. We developed a multi-step pipeline based on bash, PHP, and Python scripting languages. A sample workflow using maize as a query proteome is shown in Figure 1. Each step requires minimal configuration. As input, the pipeline uses a set of protein structures as the query proteome and one or more sets of protein structures as the target proteomes. The FASSO pipeline is optimized to predict structural orthologs for a pair of proteomes (ranging up to 60,000 proteins each) using a high-performance computing environment within 1 to 2 days. The approach infers functional annotations based on ortholog assignments. The quality of each ortholog pair is categorized based on the level of consensus of predictions from the three methods. FASSO generates Venn diagrams showing protein counts and coverage percentages between the platinum, gold, and silver ortholog predictions for each set of orthologs. The pipeline provides intermediate output files for each method, including top ten hits, reciprocal best hits, and a heat map comparing the three approaches.

**Figure 1:**
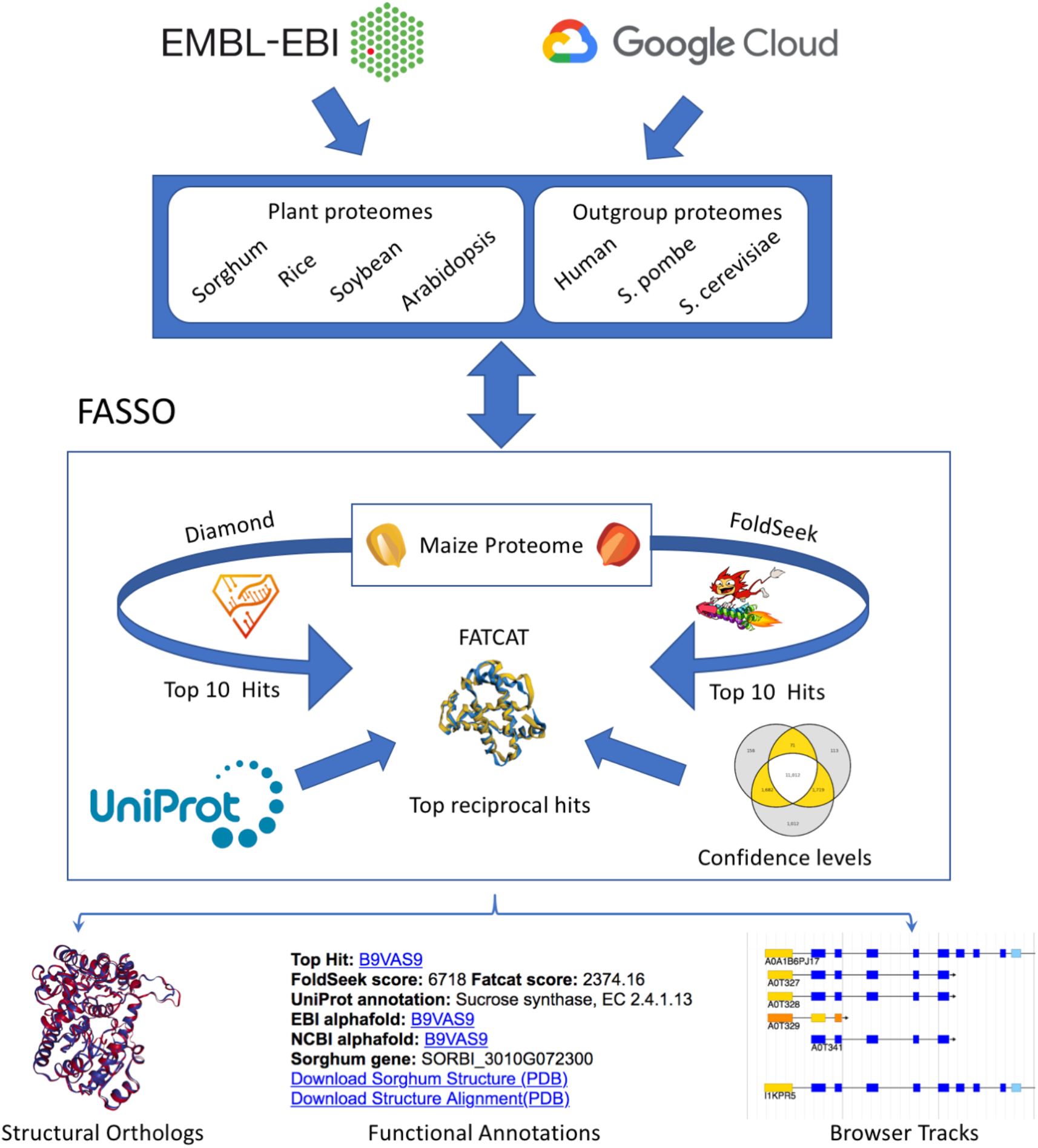
FASSO pipeline to annotate the maize proteome. The schematic diagram shows the major steps of annotating the maize proteome using FASSO. The proteomes of four plants (sorghum, rice, soybean, Arabidopsis) and three outgroups (human, *S*.*cerevisiae*, and *S. pombe*) are downloaded from the EBI AlphaFold Structure Database and Google Cloud. Each protein in the maize proteome is aligned to every protein in each target proteome using the sequence-based method Diamond and the structure-based method FoldSeek. The top ten hits from Diamond and FoldSeek are used as candidate proteins for a second sequence-based method FATCAT. The top ten RBH hits from Diamond, FoldSeek, and FATCAT are aggregated to create a final set of structural orthologs. FASSO integrates UniProt annotations to infer functional annotations and assigns confidence levels to each annotation based on the level of agreement between the three methods. The results can be integrated in other tools to visualize protein structures, curate function, or view within a genome.

The final FASSO output provides the following:

- A set of structural orthologs with confidence assignments
- Functional annotations for each protein
- A working set of flagged orthologs

FASSO was benchmarked on Ceres, a USDA-ARS funded high-performance computing infrastructure. Each pipeline step used one node, using five cores with Intel Xeon Processors (6140 2.30GHz 25MB Cache or 6240 2.60GHz 25MB Cache), and 32 GB of allocated DDR3 Memory. The DIAMOND, FoldSeek, and FATCAT steps split the target proteome into 10 to 30 subsets of proteins, and aligned each subset in parallel using a single core each.

We describe each step of the pipeline below.

### Data preparation

The FASSO pipeline requires at least two proteomes with PDB-formatted structures. The pipeline is designed for UniProt data since a large set of AlphaFold structures and functional annotations are available from UniProt. The protein structure data for this project is available from the AlphaFold Protein Structure Database (https://AlphaFold.ebi.ac.uk/) or Google Cloud. If protein structures are unavailable from these sources, FASSO can use structural annotations that are experimentally determined or computationally predicted from AlphaFold, or other prediction methods [28–30] that provide PDB results. Protein sequences are exacted from the PDB files using the program pdb2fasta (https://github.com/kad-ecoli/pdb2fasta) and stored as FASTA files. Functional annotations are downloaded in tab-delimited format from UniProt Knowledgebase (https://www.uniprot.org/uniprotkb) [40] by filtering by taxonomy and selecting the columns “Entry” and “Protein Name” for both reviewed and unreviewed annotations. A data preparation script is provided to unzip the data, create the directory structure, format the data, and build indexes and databases for future steps. The preparation step only needs to be executed once for each proteome.

### Reciprocal Best Hit (RBH) approach (Diamond)

A generic reciprocal best hit approach was used to identify sequence-based orthologs. The program Diamond (version 2.0.6) [11] was used based on its balance of speed and accuracy [13]. Each protein in the query proteome is aligned with all proteins in the target proteome (all vs. all approach). Conversely, each protein in the target proteome is aligned with the query proteome.

Here is a sample Diamond command: “diamond blastp -d TargetProteome -q QueryProteome-b12 -c1 -o ouputfile.txt” (without the quotes). Each protein’s top ten hits (based on alignment score) are saved. A RBH is identified when two proteins, one from each genome, are each other’s top-scoring hit based on the highest alignment score.

### Reciprocal Best Structure Hit (RBSH) approach (FoldSeek)

A reciprocal best structure hit approach (RBSH) [21] was developed to identify structure-based orthologs. Analogous to the RBH approach, the RBSH approach aims to identify protein pairs from different proteomes with top-scoring structural alignments. If two proteins are the top structural alignments with each other, they are identified as structural orthologs. We use FoldSeek (https://github.com/steineggerlab/foldseek) to structurally align the proteins between the proteomes. FoldSeek is many times faster than other approaches and can compare two proteomes in less than a day when using parallelization on an HPC. A sample FoldSeek command is “foldseek easy-search QueryProteome TargetDatabase output.m8 tmpfile” (without the quotes). Each protein’s top ten hits and reciprocal best structure hits (based on the highest template modeling (TM) score-a metric to measure the global similarity between two proteins) are saved.

### Reciprocal Best Structure Hit (RBSH) approach (FATCAT)

The second round of RBSH uses FATCAT (Version 2.0-https://github.com/GodzikLab/FATCAT-dist). FATCAT differs from FoldSeek by allowing twists around pivot points in the structures. FATCAT uses dynamic programming to chain together aligned fragment pairs (AFP) between the two proteins. Rotations and translations are allowed to improve the alignment of the fragment pairs. The alignments are chosen by maximizing longer AFPs and minimizing high root mean square deviations (RMSD) between the compared structures. A chaining score is calculated from the mismatched regions and gaps created by the chained AFPs [34]. The method is resource intensive, and it takes several hours to align a single protein against a proteome using an HPC environment. To overcome this resource demand, we only used FATCAT to align protein pairs from the top ten results of the Diamond and FoldSeek methods. The Perl script “FATCATQue.pl (with parameters -i ./ -m -t -ac)” (without quotes) facilitates running alignments in batch at a proteome scale. The results are sorted based on the FATCAT highest chaining scores based on the quality of the alignments and composition of extensions, gaps, and twists. The top ten hits and reciprocal best structure hits for FATCAT are saved.

### Final structural orthologs and functional annotations

The final step of FASSO combines the reciprocal best hits from Diamond, FoldSeek, and FATCAT. Each ortholog receives a confidence label of either platinum, gold, or silver based on the consensus between the three methods. If all methods predict the same ortholog pair, it is labeled platinum; if two of the three methods agree, it is labeled gold; and if only a single approach makes the prediction, it is labeled silver. If conflicting predictions occur for a protein in the query proteome, they are removed from the ortholog set and added to a “working set” of orthologs. Functional annotations from the UniProt Knowledgebase are assigned to each query protein based on annotations of the target protein in the ortholog pair. FASSO merges results into a single annotation file if multiple target proteomes are used. For downstream analysis, the final ortholog predictions for the maize genome were aligned back to the B73 maize reference genome using the miniprot [41] command “miniprot -ut16 --gff maize_database.mpi target_pdb.fa > output.gff” (without quotes). The query database was built using the genomic coordinates of Zm-B73-REFERENCE-NAM-5.0_Zm00001eb.1 [42].

## Results and discussion

### AlphaFold database

We used FASSO to predict structural orthologs in maize from the plant proteomes retrieved from the AlphaFold Protein Structure Database. The AlphaFold Protein Structure Database provides bulk downloads for 48 species including four plant model organism proteomes: *Arabidopsis thaliana* (Arabidopsis), *Glycine max* (soybean), *Oryza sativa* (Asian rice), and *Zea mays* (maize). A fifth plant species, *Sorghum bicolor*, (sorghum) was downloaded from the Deepmind/AlphaFold public dataset on the Google Cloud using taxonomy id 4558. Sorghum is an important crop and close relative to maize (both belong to the Andropogoneae tribe) [43]. We chose three well-annotated proteomes as outgroups: Homo sapiens (human), *Saccharomyces cerevisiae* (*S. cerevisiae*, budding yeast), and *Schizosaccharomyces pombe* (*S. pombe*, fission yeast). The protein structure models for all the proteomes are from the AlphaFold Database Release 4 (July 2022). The plant proteomes and outgroups we used are diverse in several ways (Table 1). First, the eight species represent a broad set of taxonomic families with a focus on plants and crops. Second, the number of proteins in each proteome ranged from 5,128 for *S. pombe* to 55,799 for soybean. Third, the prediction structure quality distribution was different across the proteomes.

**Table 1:** AlphaFold proteome summary and structure prediction quality. The table lists the UniProt identifier, taxonomy family, protein count, residue quality ratio, and average residue scores for each of the eight proteomes. Proteomes are listed in alphabetical order.

| Proteome | UniProt | Family | Number of proteins | Residue quality ratio | Average residue score |
| --- | --- | --- | --- | --- | --- |
| Arabidopsis | UP000006548 | Brassicaceae | 27,434 | 2.2 | 76.2 |
| Human | UP000005640 | Hominidae | 23,391 | 1.6 | 74.3 |
| Maize | UP000007305 | Poaceae | 39,299 | 1.6 | 71.6 |
| Rice | UP000059680 | Poaceae | 43,648 | 1.4 | 68.1 |
| Sorghum | UP000000768 | Poaceae | 42,553 | 1.5 | 68.9 |
| *S. cerevisiae* | UP000002311 | Schizosaccharo mycetaceae | 6,040 | 2.2 | 77.1 |
| *S. pombe* | UP000002485 | Schizosaccharo mycetaceae | 5,128 | 2.4 | 78.8 |
| Soybean | UP000008827 | Fabaceae | 55,799 | 1.9 | 74.8 |

### Quality of FASSO Ortholog Prediction

Figure 2 shows the relationship between the average structure quality (x-axis) and the alignment score of FATCAT (y-axis) between maize and sorghum; each point is color-coded by the quality of the ortholog prediction. The quality of the alignments increases with the quality of the structures. The higher-quality structures were also more likely to provide platinum-quality orthologs. We focused on the maize genome as our reference use case for a few key reasons: (1) maize is an important agronomic crop and scientific model organism [44, 45]; (2) maize is a model plant species with a wealth of genetic, genomic, and phenomic datasets and resources [42, 46]; (3) maize is well studied with a few hundred well-annotated genes [47, 48], but has limited experimentally determined functional annotations at the genome-scale; and (4) there are annotated genomes at multiple levels of taxonomic distances from maize (Figure 3 shows the phylogeny relationship between the eight species built with the ETE Toolkit [49] using NCBI Taxonomy database [50]).

### AlphaFold Protein Structure Prediction Quality

We ran FASSO for each pairwise combination of the eight selected genomes (full results in Supplementary Datasets #1 and #2). Functional annotations were merged into a single file (Supplementary Dataset #3). We provide three metrics to describe the quality of the protein structure predictions at a proteome scale. The first metric is the residue quality ratio. We calculated this ratio by dividing the number of residues within a proteome with a high or very high confidence score (≥ 70 pLDDT) by the number of residues with a low or very low confidence score (< 70 pLDDT). We chose ratios greater than 2.0 as a cutoff score, as the ratio represents when there are at least twice as many high-confidence residues as low-confidence residues. This ratio was below 2.0 for human, maize, rice, sorghum, and soybean and above 2.0 for Arabidopsis, *S. cerevisiae*, and *S. pombe*. The second metric is the average residue score across all the proteins in each proteome. The average residue score was below 70 pLDDT for rice (68.1) and sorghum (68.9); the highest score was 78.8 for *S. pombe*. The third metric is the distribution of the average per-residue score per protein. Figure 4 shows a plot of these distributions. Each of the proteomes had a peak near an average pLDDT score of 90, but the three outgroups, human and the two yeast genomes, had much higher and steeper peaks at this value. Each of the five plant proteomes had a second peak between 55 and 65 pLDDT, with the three grass (Poaceae) proteomes (maize, sorghum, and rice) with the most prominent bi-modal distributions. The three grass genomes also had the highest number of proteins with an average pLDDT labeled as low or very low (< 50 pLDDT). Together, these metrics show that the three outgroups had a more uniform quality of high-confidence predictions than the plant genomes. The smaller proteomes with less complex genomes, Arabidopsis and the two yeast proteomes, have the highest residue quality ratio compared to the larger and more complex genomes (e.g, whole-genome duplications events and copy-number variations). These distinctions in structural quality are important to note as they reflect on either the quality of the protein structure predictions or gene structure annotations, which impact predicting structural orthologs.

**Figure 2:**
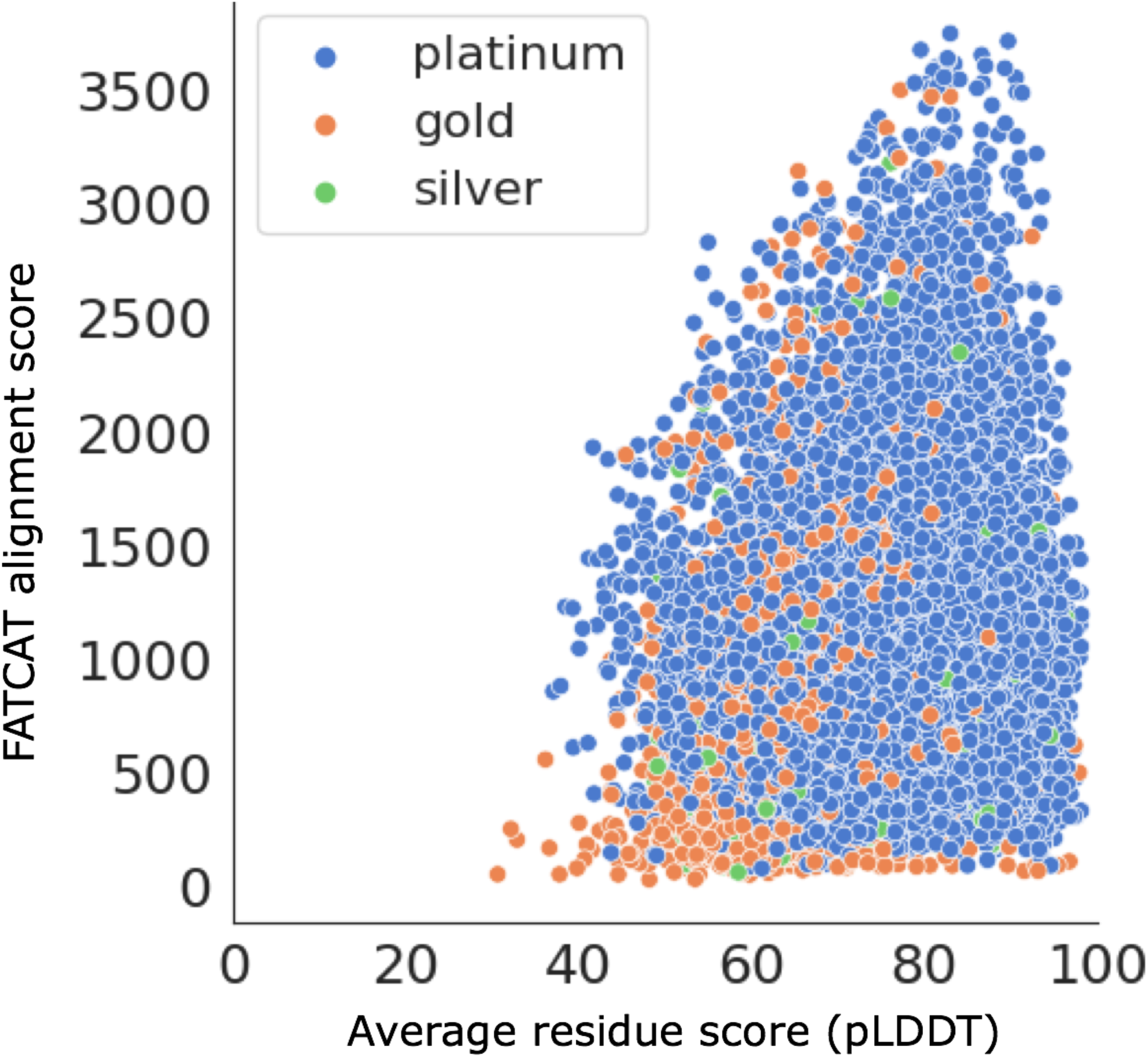
A scatter plot showing the relationship between a protein’s quality and alignment scores for FASSO-predicted orthologs between maize and sorghum. The protein structure quality (x-axis) is calculated by taking the average of all the residue scores (pLDDT-which ranges from 0 (low) to 100 (high)) for each maize protein. The y-axis represents the FATCAT chaining scores between the maize protein and the FASSO ortholog prediction from sorghum. Each point is color-coded by the type of ortholog prediction. The quality of the alignments increases with the quality of the structures.

**Figure 3:**
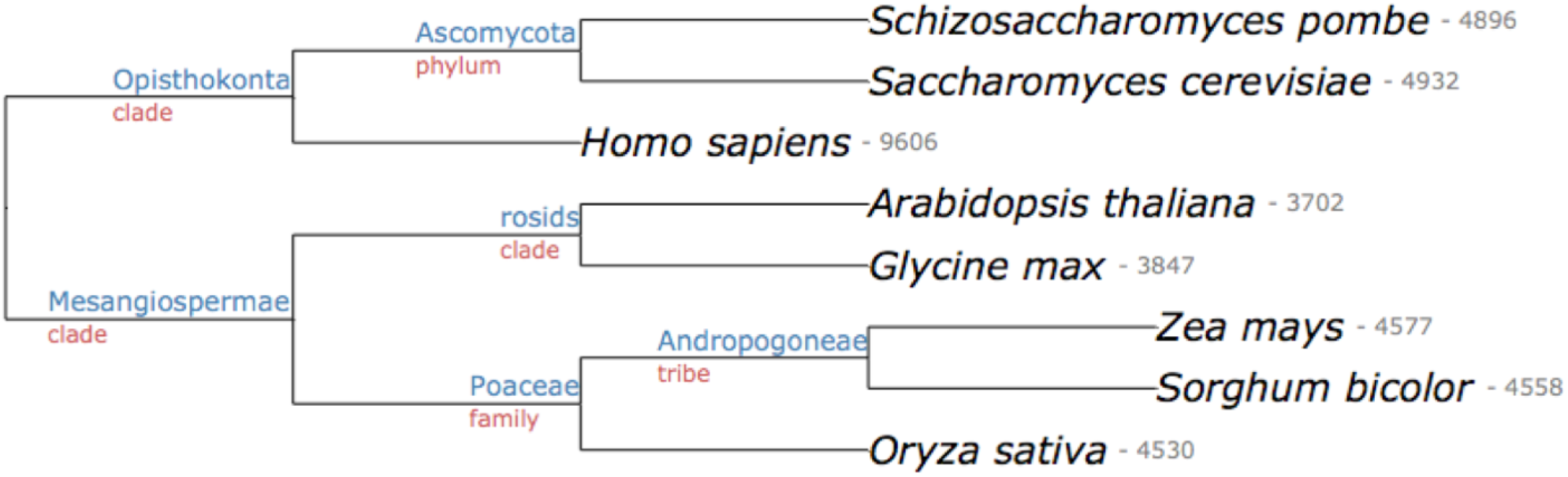
Phylogenetic tree for each of the eight genomes. This study benchmarked FASSO using proteomes from five plant species and three distant outgroups. The eight species are shown on the phylogenetic tree. The number next to the scientific name is the NCBI Taxon identifier. Each junction of the tree lists the common tribe, family, or clade to which the species belong. The tree was built using the ETE Toolkit and the NCBI Taxonomy database.

**Figure 4:**
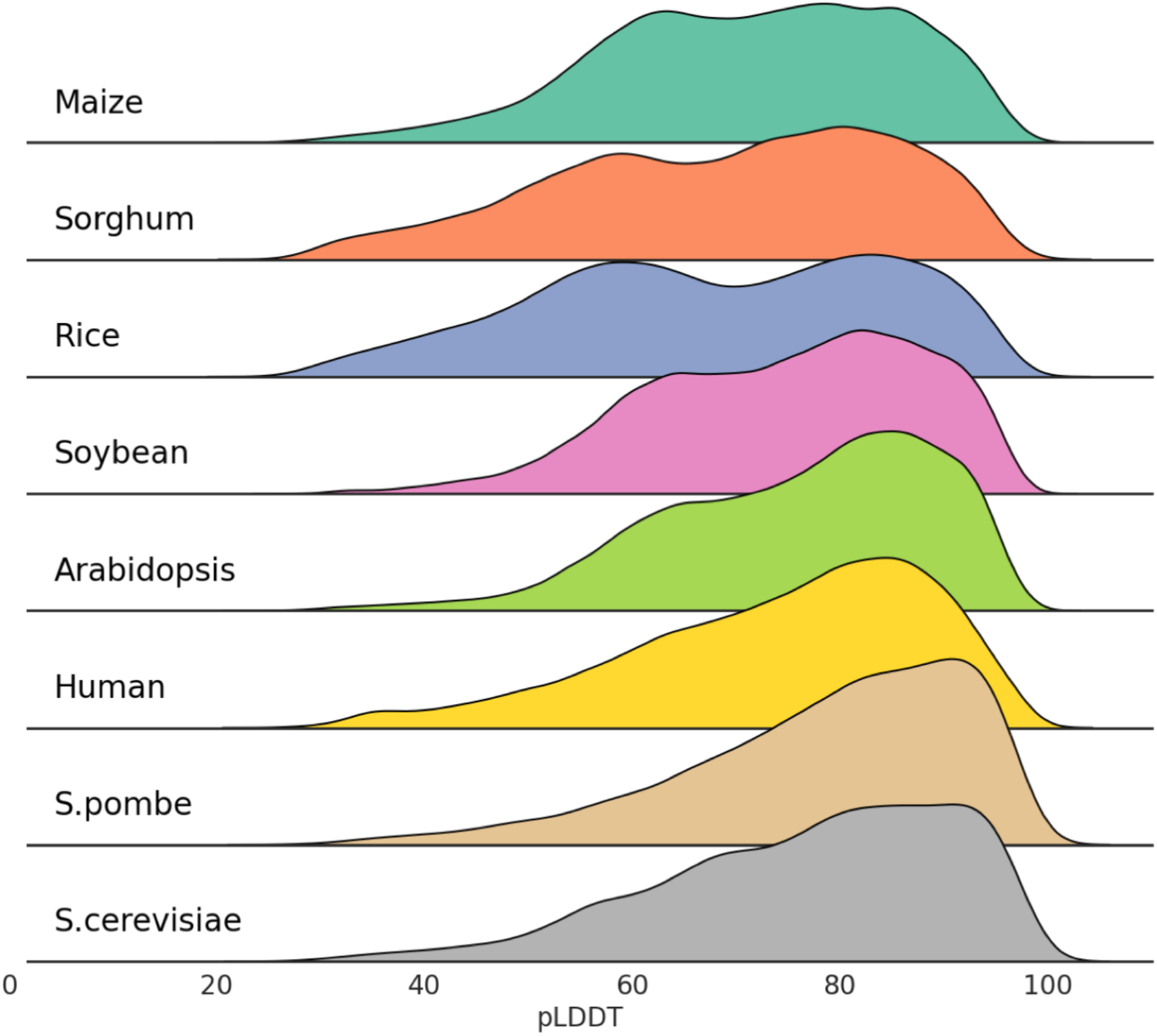
Ridgeline plot of the distribution of average residue score per protein across each of the eight proteomes. Each residue in an AlphaFold structure has an average residue score (pLDDT) ranging from 0 to 100. Each protein in the eight proteomes is assigned a confidence score by averaging the pLDDT for each residue in the protein. The ridgeline plot shows the distributions of these scores for each of the eight proteomes.

### Performance Comparison of Diamond, FoldSeek, and FATCAT

As described, the first step of the FASSO pipeline identifies potential orthologs between proteomes using three existing approaches: Diamond, FoldSeek, and FATCAT. To compare the results, we used each method’s top-ten ranked hits sorted first by the method’s respective score, then by either e-value or p-value (Supplementary Dataset #4). The alignment score provided better results than the e-value or p-value since it is independent of the size of the proteomes and was less likely to have hits with identical scores (e.g., multiple e-values of 0.0). Both Diamond and FoldSeek can make all-vs-all comparisons between proteomes in a practical timeframe. For example, using an all-vs-all approach between the maize and rice proteomes requires over 1.7 billion comparisons. On an HPC, Diamond finished the all-vs-all run in a few minutes, and FoldSeek completed its run in less than a day. The drawback is that FATCAT can take up to a day to run a single protein against a proteome, so running an all-vs-all approach is not practical. For this reason, we only run FATCAT alignments with the merged top ten predictions from Diamond and FoldSeek, which has a 300-fold to 6,000-fold reduction in the number of protein alignments. Each alignment has a FATCAT score that can be used for a consistent metric to benchmark results. To compare the predictions for the three methods, we used two types of evaluation: how well each method’s predictions agreed with each other (agreement) and the number of proteins in the query proteome with a prediction ortholog (coverage).

To measure the level of agreement between methods, we first compared the top ten ranked hits between Diamond and FoldSeek. Figure 5A compares these two methods using maize as the query proteome and sorghum as the target proteome. For each maize protein, up to ten hits are reported for each method and ranked based on score. The rows of the table are labeled #1-#10 by Diamond rank, and the columns are labeled #1-#10 by FoldSeek rank. The “no hit” row and column display the count of proteins that are ranked by one method, but are either not predicted or ranked below the top ten for the other method. Each cell of the heat map is colored by the counts of proteins that have that specific combination of ranks. For example, the upper left cell has 27,182 proteins with the same top-ranked (#1) sorghum protein hit in both Diamond and FoldSeek. This is significant since these orthologs were identified as the top hit for both methods. As expected, the largest number of the predictions agreed with each other (the cells in the diagonal of the matrix), but thousands of predicted hits disagreed. This is especially true for proteins labeled “no hit”, which corresponds to the bottom row and the column furthest to the right. These are the proteins in which additional structural information can provide improved ortholog annotation predictions.

**Figure 5:**
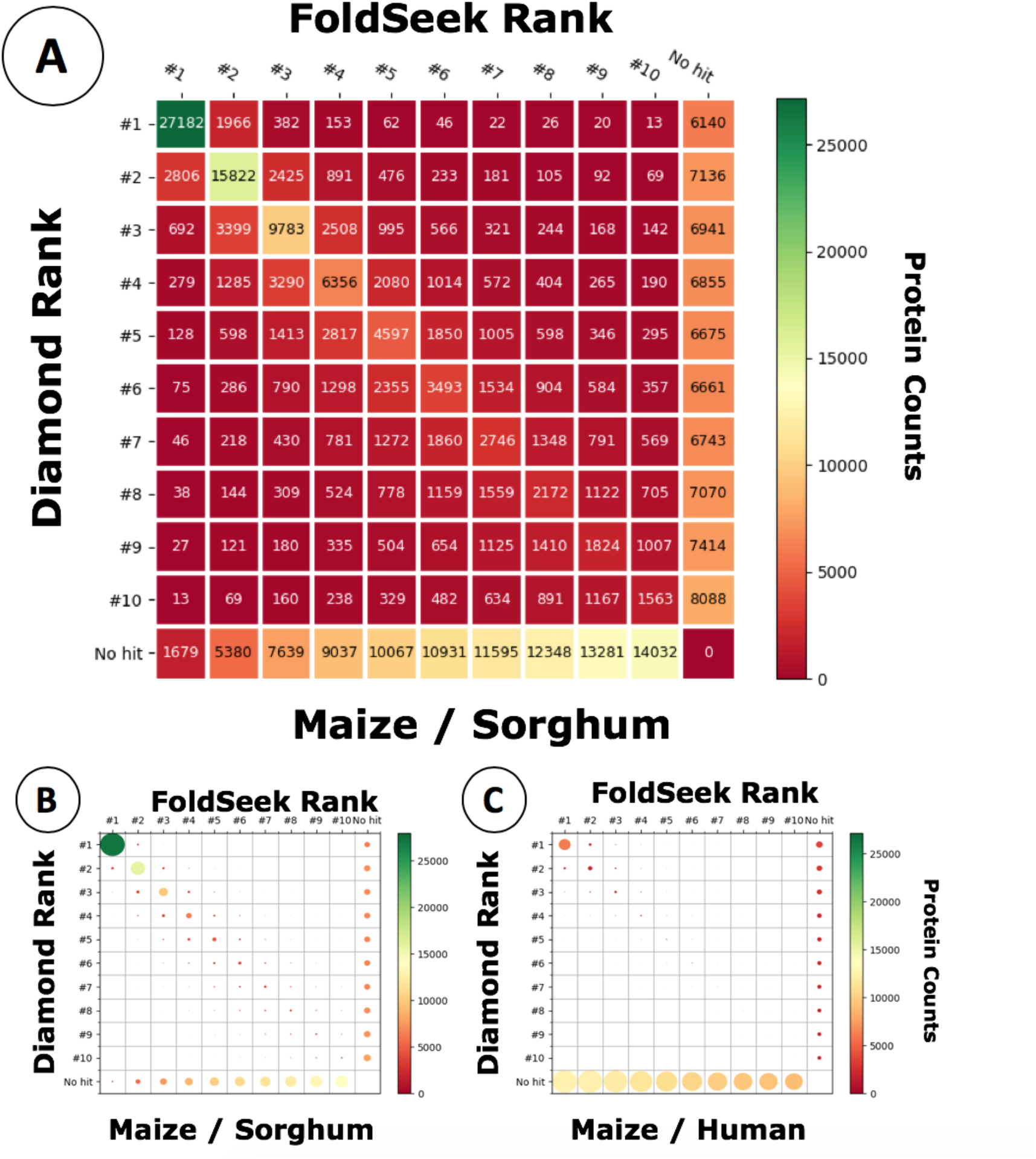
Heatmaps of Diamond and FoldSeek ranked hits. Panel A shows a heatmap of protein counts based on Diamond and FoldSeek ranked hits between maize and sorghum. The heatmap is color-coded by protein counts. Each column represents the proteins with a specific FoldSeek rank based on alignment score from #1 (top hit) to #10 (10th best hit). Each row represents the proteins with a specific Diamond ranking based on alignment score from #1 to #10. Each square is the count of proteins having both rankings from the two methods. Panel B and C show similar data comparing the ranked hits between maize and sorghum (panel B) and between maize and human (panel C). The count of proteins determines the color of each circle based on the global minimum and maximum of the overall protein count distribution across the two pairs of proteomes (e.g. maize/sorghum and maize/human), and the size of each circle is based on the local minimum and maximum of either pair of proteomes (e.g. maize/sorghum).

Figure 5B and 5C compare the results of Diamond and FoldSeek between closely related species (Panel B: maize/sorghum) and distantly related species (Panel C: maize/human). A complete set of heat maps are found in Supplementary Dataset #5. The count of proteins determines the color of each circle based on the distribution of protein counts across the two pairs of proteomes, maize/sorghum and maize/human. The size of each circle is based on the distribution of protein counts between maize/sorghum in Panel B and maize/human in Panel C. The maize/sorghum results have a higher proportion of hits in an agreement between the two methods. The maize/human results have a higher proportion of predictions absent (i.e. “no hit”) in either Diamond or FoldSeek. This is especially true for FoldSeek predictions. For example, there were over twice as many top-ranked FoldSeek predictions not found by Diamond (12,534) as there were top-ranked predictions that were also top-ranked in Diamond (6,054). Similar trends are seen across all the other species where agreement is high between more closely related species, and FoldSeek identified large sets of predictions not found in Diamond when the species are distantly related (see Supplementary Dataset #5 for all comparisons between Diamond, FoldSeek, and FATCAT). These findings support the notion that sequence similarity (and therefore sequence homology sensitivity) diverges faster than structure similarity [17, 18, 51] between orthologs, showing the importance of using structure to verify existing orthologs or annotate novel orthologs.

In addition to quantifying how often the methods predicted similar results, coverage measures the number of proteins in a proteome with a predicted ortholog. Figure 6 presents heat maps that compare percent agreement and coverage for each pairwise combination of proteomes. Figure panels 6A and 6B show two heat maps comparing the level of agreement between Diamond /FATCAT and FoldSeek/FATCAT. The plots only show the proportion of hits where the two methods had the same top-ranked hit. Diamond and FoldSeek predicted over 50% agreement for most proteome combinations, with the highest percentage between the grasses (maize, sorghum, and rice) and between the three non-plants (human, *S. pombe*, and *S*.*cerevisea*). Although Diamond is sequence-based and FoldSeek is structure-based, Diamond agrees with FATCAT better than FoldSeek in some instances, including when soybean, human, and the two yeast genomes were the target proteome. Diamond, FoldSeek, and FATCAT predictions often disagreed when using the highly duplicated soybean genome as a target proteome. Up to 75% of the soybean genes have multiple copies [52], causing a high level of one-to-many or many-to-many orthology relationships between a query proteome and the soybean proteome. The RBH approach only identifies the best one-to-one orthology relationship to maintain higher confidence at the sacrifice of a lower proteome coverage [3]. If the protein sequences from the gene copies are very similar, it is more likely that the different RBH approaches may identify conflicting orthologs. Similar to the soybean genome, maize has gone through two whole genome duplication events, but the maize genome has higher rates of gene loss and had more distinct subgenomes before polyploidization [53]. Therefore, the sequence and structure-based methods agreed more often for gene pairs in the maize proteome than soybean. The percent agreement between Diamond/FATCAT and FoldSeek/Diamond was nearly the same when using either maize or sorghum as the target proteome, even though maize has had a whole genome duplication event after its divergence from sorghum.

**Figure 6:**
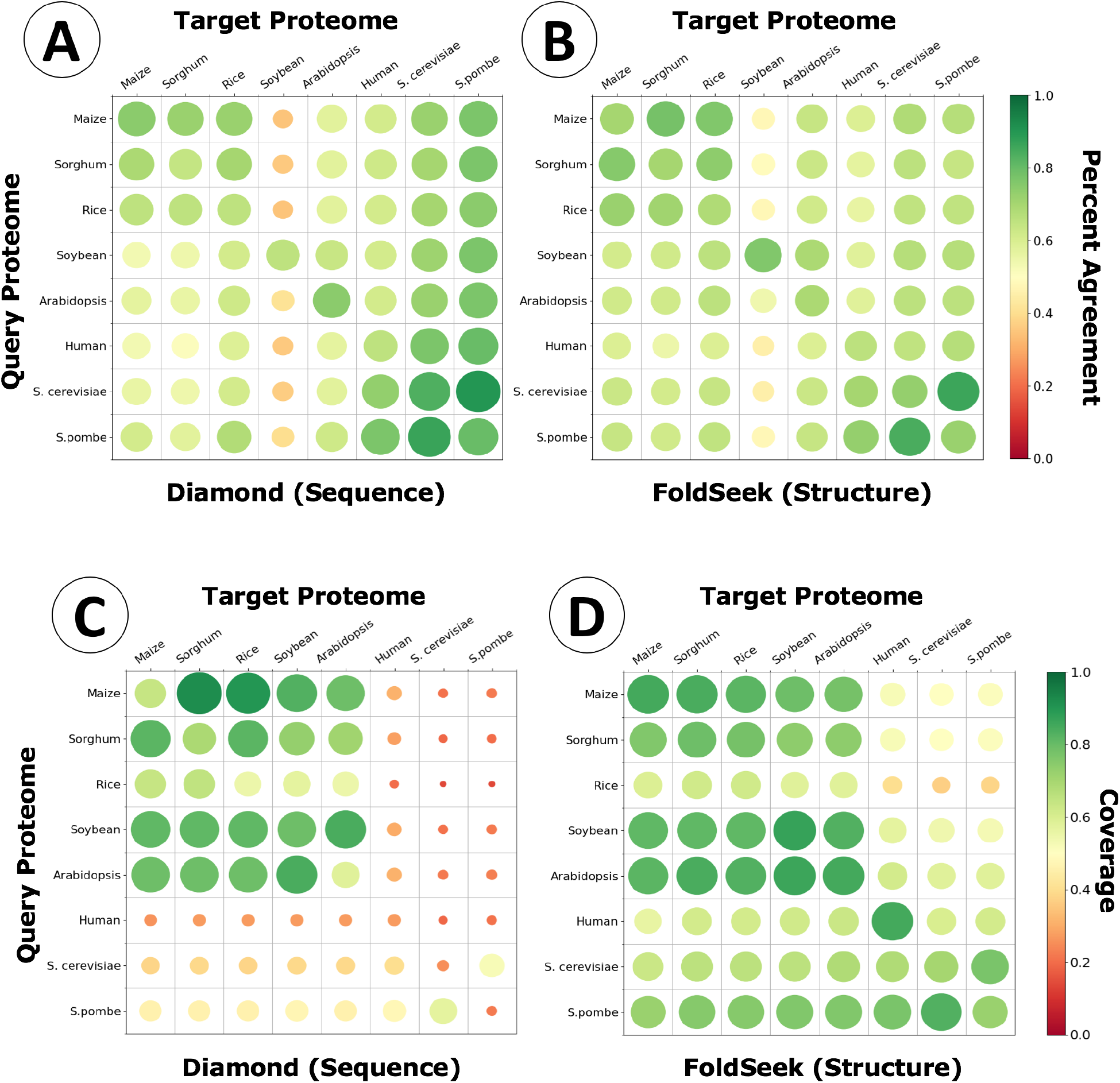
Agreement and coverage heatmaps. Panels A and B show heatmaps where the size and color of each circle are determined by the percentage of top-ranked Diamond (panel A) or FoldSeek (panel B) predictions that agree (agreement) with the top-ranked FATCAT predictions. The rows list the query proteome and the columns list the target proteome. Panels C and D show heatmaps where the size and color of each circle are determined by the percentage of the query proteome (coverage) that has at least one top-ranked prediction from Diamond (panel C) or FoldSeek (panel D). The rows list the query proteome and the columns list the target proteome.

The two additional heat maps provided in Figures 5C and 5D compare the number of proteins within a proteome (coverage levels) with a predicted ortholog of each method. We compare the sequence-based approach Diamond to the structure-based FoldSeek. We did not directly report the coverage results of FATCAT since those results are dependent on the predictions of Diamond and FoldSeek. The differences in coverage between the two methods are more prominent than the differences in agreement. Diamond has 50% or greater coverage for 27 proteome pairs, including all combinations of plant proteomes and greater than 80% coverage for 10 proteome pairs, but has lower coverage (< 40%) for 36 pairs of proteomes and very low coverage (<20%) for 15 proteome pairs. FoldSeek had 50% or greater coverage for all but three proteome pairs (rice/human, rice/*S. cerevisiae*, and rice/*S. pombe*), and greater than 80% coverage for 15 proteome pairs. The structure-based predictions provide as many or, in most cases, a larger set of potential orthologs across proteomes.

### FASSO and Reciprocal Best Hit results for ortholog predictions

The next major step of FASSO uses Diamond, FoldSeek, and FATCAT to find reciprocal best hits and aggregates those results for a final set of ortholog predictions. FASSO merges the results from each method, assigns confidence labels based on the level agreement (see Methods), and removes conflicting predictions. Our approach increases the number of high-quality predictions and removes potential erroneous annotations identified from the individual methods. Although using each method in a one-vs-all approach is straightforward and provides high coverage, it suffers from several setbacks [54]. The main concern is that gene duplication events can cause two or more proteins in one proteome to have high sequence similarity but with different functions. This is especially true in plant species where gene duplication events are more common. A second concern occurs when a high sequence or structure similarity exists at local regions across multiple proteins, but the global similarity could be quite different. The reciprocal best hit approach partially solves this problem by providing bi-directional support for a prediction, but FASSO provides even stronger support by combining sequence-based and structure-based predictions.

Figure 7 shows how the sequence-based and structure-based scores are related. Figure 7A provides a scatter plot showing the relationship between the Diamond alignment score (x-axis) and the FATCAT alignment score (y-axis). Similarly, Figure 7B is a scatter plot of the FoldSeek alignment score (x-axis) and the FATCAT alignment score (y-axis). Each point is blue if it was a FASSO annotation or orange if it was removed for conflicting predictions (working set). Both plots show a strong correlation between each method’s scores and FASSO predictions tend to have overall high structure scores compared to the working set. The Pearson correlation coefficient is 0.82 between Diamond and FATCAT scores and 0.92 between FoldSeek and FATCAT scores. A stronger correlation was expected between the two structure-based approaches. The plots show many examples where the FATCAT score was higher relative to either the Diamond or FoldSeek score, but few examples where the FATCAT score was relatively lower than Diamond or FoldSeek. One example (highlighted with red arrows) shows a predicted ortholog pair (A0A1P8B4E4 and O60055) with a low sequence alignment score (Diamond score 323.9) and relatively high structure alignment scores (FoldSeek score: 4567 and FATCAT score: 5164.44). Figure 7C shows the FATCAT structure alignment between the AlphaFold structures of the two proteins. Although the sequence similarity was only 20% with 12% of the alignment as gaps, the structures were still well-conserved.

**Figure 7:**
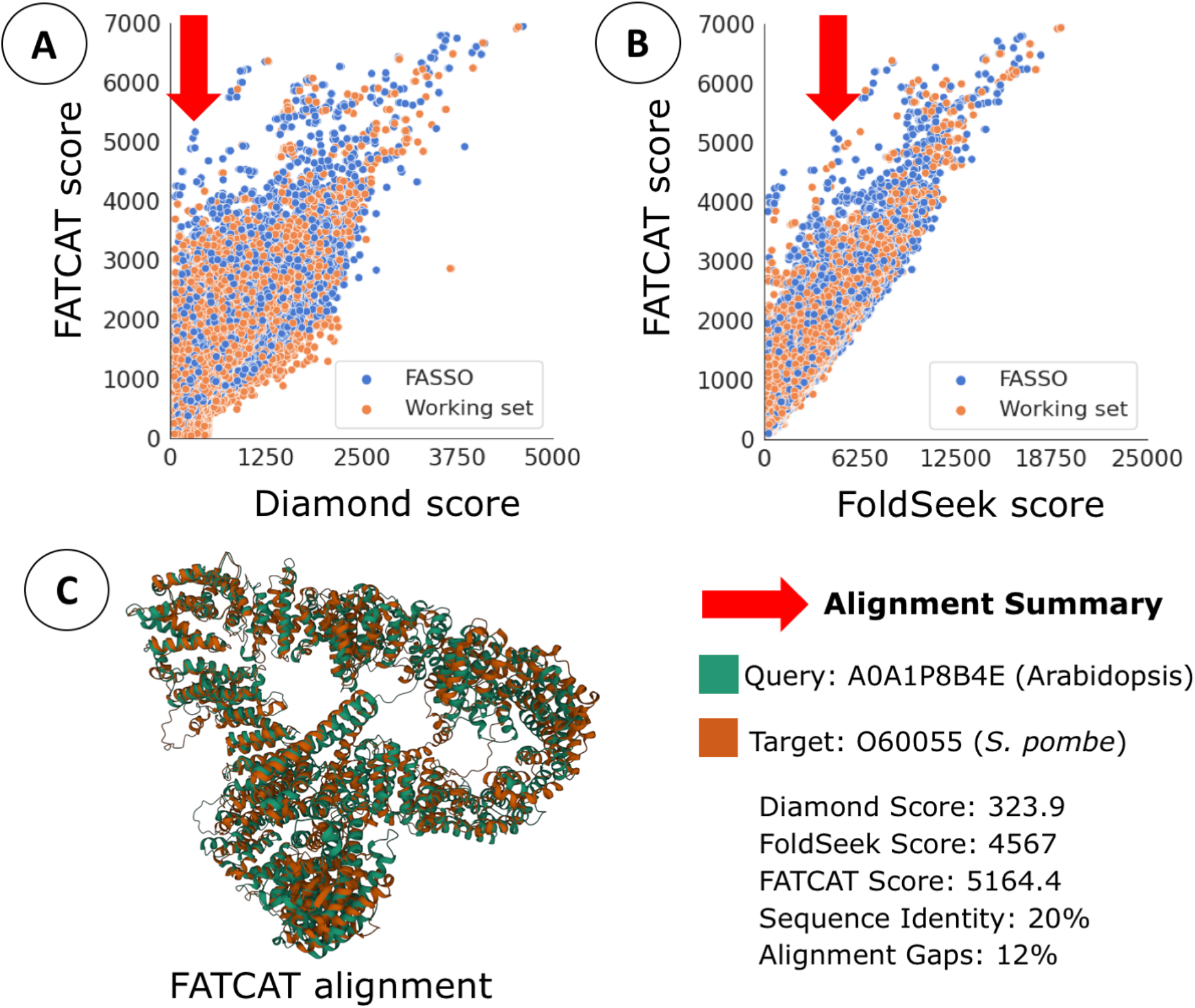
Scatter plot of the relationships between Diamond/FATCAT and FoldSeek/FATCAT alignment scores. Panel A shows the relationship between the Diamond alignment score (x-axis) and the FATCAT chaining score (y-axis). Panel B shows the relationship between the FoldSeek TM score (x-axis) and the FATCAT chaining score (y-axis). Each point on the two panels is either a FASSO-predicted ortholog (blue) or an ortholog in the working set (orange). The red arrow highlights an example where the sequence alignment score was low and the structure alignment score was high. This example is a FASSO-predicted ortholog between the proteins A0A1P8B4E (Arabidopsis) and O60055 (*S. pombe*). Panel C displays the composite 3D structural alignment between A0A1P8B4E (green) and O60055 (brown)Details about the alignment of the three methods for this ortholog pair are provided next to the structure alignment.

We ran FASSO on all pairwise combinations between eight species and compared the 64 pairs of proteomes using over 170 billion protein alignments. Each proteome was also compared with itself (after removing self-identifying protein hits) to identify a set of homeologs. Figure 8 shows the coverage of the four approaches for each query proteome against the target proteomes. The target proteomes are listed in increasing order based on the estimated years of species divergence. Some trends emerge from these results. Proteome coverage dropped by as much as 40-50% when using the reciprocal-based approaches compared to using only best hits. Each method predicts an ortholog for 20-50% of the proteins when comparing two plant proteomes (see Supplementary Datasets #1 and #2 for exact counts). Overall proteome coverage decreased (below 10% for the outgroups) as the evolutionary distance increased. FATCAT predictions provide the highest coverage when comparing plant proteomes, but had lower coverage than both Diamond and FoldSeek for the outgroups. The three smallest proteomes (Arabidopsis, *S. cerevisiae*, and *S. pombe*) have the highest overall coverage. The proteomes of human (with no closely related species) and soybean have the lowest coverage. Soybean even has low coverage with other plants, likely due to the limitation of the RBH method with a high number of gene copies [55]. When using the plant proteomes as the query and the outgroup proteomes as the target, the coverage dropped below 10%. FASSO has the lowest coverage by design as ortholog predictions with conflicting results are removed.

**Figure 8:**
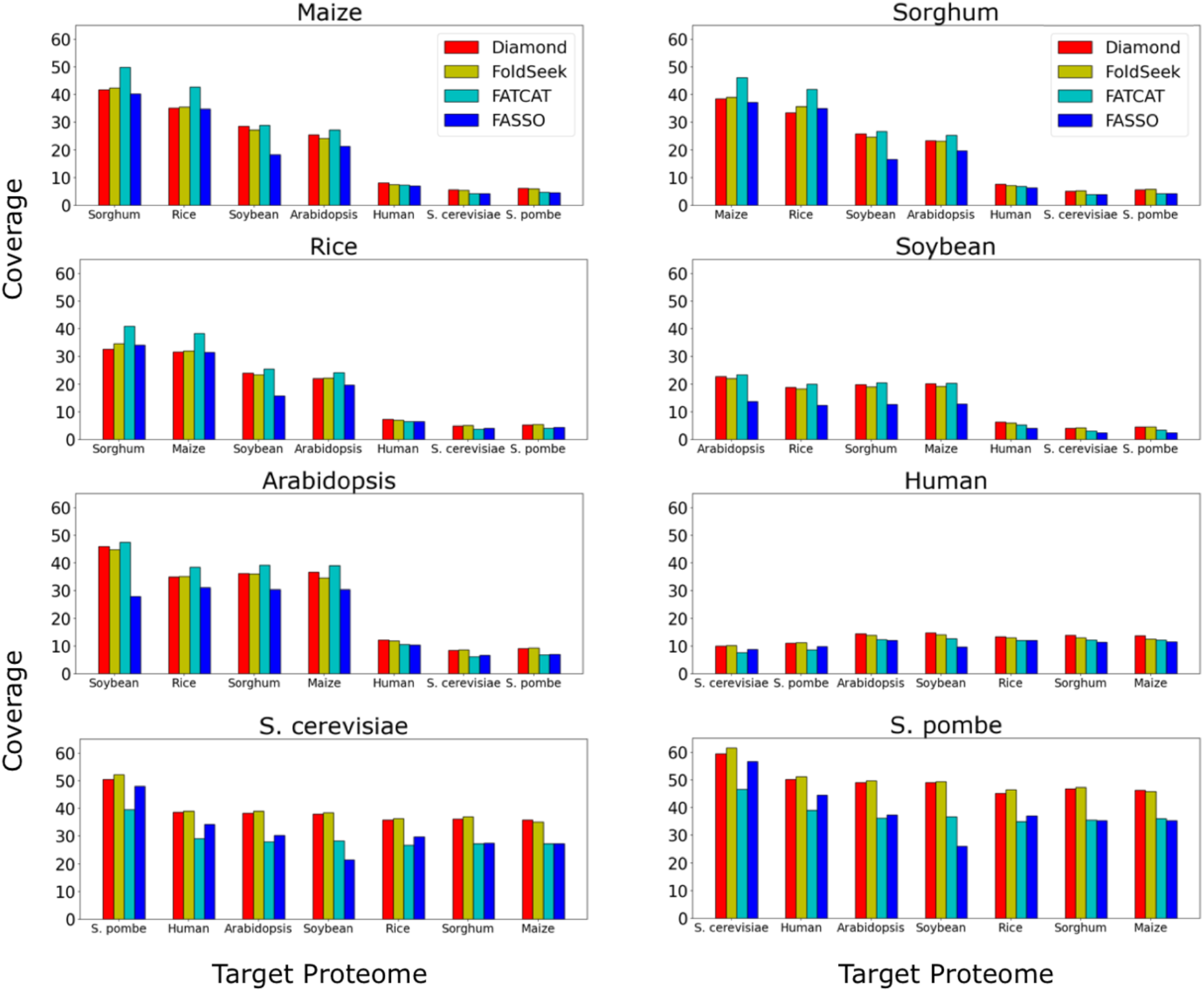
Bar charts of annotation coverage. Eight bar charts show results using each species as a target proteome (labeled on the top of each chart) with the other seven proteomes (x-axis). Each chart lists species in ascending order based on the time the proteome diverged from a common ancestry with the query species. The y-axis represents the percent coverage of the target proteome with a predicted ortholog using FASSO, and the reciprocal best hit results from Diamond, FoldSeek, and FATCAT.

### Use-case: Annotating the maize proteome

The final step of the FASSO pipeline assigns functional annotations from UniProt to the query proteome. Diamond, FoldSeek, and FATCAT assigned 2.1 million functional annotations to the 243,292 proteins in the eight proteomes. FASSO combined these results and removed conflicting predictions to produce a final set of 271,278 functional annotations (see Table 2). Supplementary Dataset #3 contains the complete annotation tables for each proteome. To illustrate how FASSO provides improved annotations over a standard RBH approach, we present a use case with the maize proteome. The maize proteome, downloaded from UniProt (Zea mays: UP000007305), has 39,299 proteins with 36,967 proteins having existing functional annotations, including 9,130 annotations with a label of “Uncharacterized protein.’”

**Table 2:** Proteome annotation summary. For each of the eight proteomes, the table lists the protein count, total counts of predicted annotations by Diamond, FoldSeek, and FATCAT, total counts of assigned annotations by FASSO, the count of uncharacterized proteins, and the count of uncharacterized proteins annotated by FASSO. Proteomes are listed in alphabetical order.

| Proteome | Protein count | Total predicted annotations | FASSO annotations | Uncharacterized protein count | Uncharacterized protein FASSO annotations |
| --- | --- | --- | --- | --- | --- |
| Arabidopsis | 27,434 | 251,704 | 39,374 | 4,675 | 1,483 |
| Human | 23,391 | 119,302 | 17,556 | 390 | 10 |
| Maize | 39,299 | 395,649 | 51,180 | 9,103 | 1,021 |
| Rice | 43,648 | 277,850 | 50,385 | 1,964 | 1,016 |
| Sorghum | 42,553 | 391,013 | 52,108 | 23,004 | 7,052 |
| *S. cerevisiae* | 6,040 | 44,434 | 13,163 | 650 | 114 |
| *S. pombe* | 5,128 | 53,857 | 13,934 | 969 | 480 |
| Soybean | 55,799 | 582,409 | 33,578 | 4,675 | 1,483 |
| Total | 243,292 | 2,116,218 | 271,278 | 45,430 | 5,614 |

Table 3 shows the number of orthologs predicted by the reciprocal best approaches from Diamond, FoldSeek, and FATCAT for the maize proteome against the seven target proteomes. The column label “New orthologs” lists both the count and percent increase of orthologs identified using the structure-based approaches (FoldSeek and FATCAT) that were not predicted by the sequence-based Diamond method. The increase in maize proteins with a structure-based ortholog is relatively small (2-3%) for the closely related grass species (sorghum and rice), but increases to a 30-40% increase in predictions for the outgroups. The table also shows the count and percent of orthologs removed when a sequence-based prediction was inconsistent with either structure-based prediction. The number of orthologs removed from the sequence-based predictions ranged from 21-53%. These results show that many proteins with the highest sequence similarity did not share the highest structural similarity and vice versa. FASSO will remove these orthologs and place them in a “working set.” Removing low-quality annotations is important since annotation errors often are propagated from one proteome to another [56]. FASSO provides a list of the working set sorted by the difference in FATCAT scores.

**Table 3:** Maize reciprocal best hits (RBH) results. The table lists the RBH results for Diamond, FoldSeek, and FATCAT when maize was the query proteome. For each of the seven target proteomes, the table lists the count of orthologs predicted by Diamond, FoldSeek, and FATCAT. The column “New Orthologs” shows both the count and percent increase of orthologs identified using the structure-based approaches (FoldSeek and FATCAT) that were not predicted by the sequence-based Diamond method. The column “Removed Orthologs” shows the count and percent of orthologs removed when a sequence-based prediction was inconsistent with either structure-based prediction. The target proteomes are listed in ascending order based on the time the proteome diverged from a common ancestry with maize.

| Proteome | Diamond-based Orthologs | FoldSeek-based Orthologs | FATCAT-based Orthologs | New Orthologs | Removed Orthologs |
| --- | --- | --- | --- | --- | --- |
| Sorghum | 19,605 | 16,629 | 16,383 | 345 (+2%) | 4,183 (-21%) |
| Rice | 16,756 | 13,963 | 13,798 | 554 (+3%) | 3,647 (-22%) |
| Soybean | 11,335 | 10,650 | 11,179 | 1,916 (+17%) | 6,060 (-53%) |
| Arabidopsis | 10,683 | 9,497 | 10,044 | 1,179 (+11%) | 3,569 (-33%) |
| Human | 2,828 | 2,932 | 3,208 | 939 (+33%) | 1,028 (-36%) |
| *S. cerevisiae* | 1,645 | 2,119 | 2,159 | 696 (+42%) | 600 (-36%) |
| *S. pombe* | 1,843 | 2,343 | 2,368 | 708 (+38%) | 685 (-37%) |
| Total | 64,695 | 58,133 | 59,139 | 6,337 (+10%) | 19,772 (-31%) |

Figure 9 presents an example of the three individual methods predicting three different orthologs when using the Arabidopsis proteome to annotate the maize query protein A0A1X7YER2 (Panel A). The protein structures and alignment scores of the top Diamond, FoldSeek, and FATCAT hits are shown in Panels B, C, and D. The protein Q9FMR1 (function label “Rho GTPase-activating protein 1 (Rho-type GTPase-activating protein 1)) has the best Diamond alignment score, but was not a top ten hit in FoldSeek. It also had the lowest FATCAT score of 392.3. The protein Q9LSL9 (with function label “Pentatricopeptide repeat-containing protein” linked to gene model At5g65560) was not identified by Diamond, but has the top alignment score in FoldSeek (3118) and a high FATCAT alignment score (2388.16). The third protein Q9LVQ5 has the same functional annotation as Q9LSL9, but was linked to a different Arabidopsis gene model: At5g55840. Q9LVQ5 was also not identified by Diamond, but has the best FATCAT score of 2516.53 and a FoldSeek score of 2390. The structures and functional annotations differed between the sequence-based and the two structure-based predictions. The FASSO method flags and moves these conflicting predictions to a working set to allow further evaluation before assigning a final ortholog assignment.

**Figure 9:**
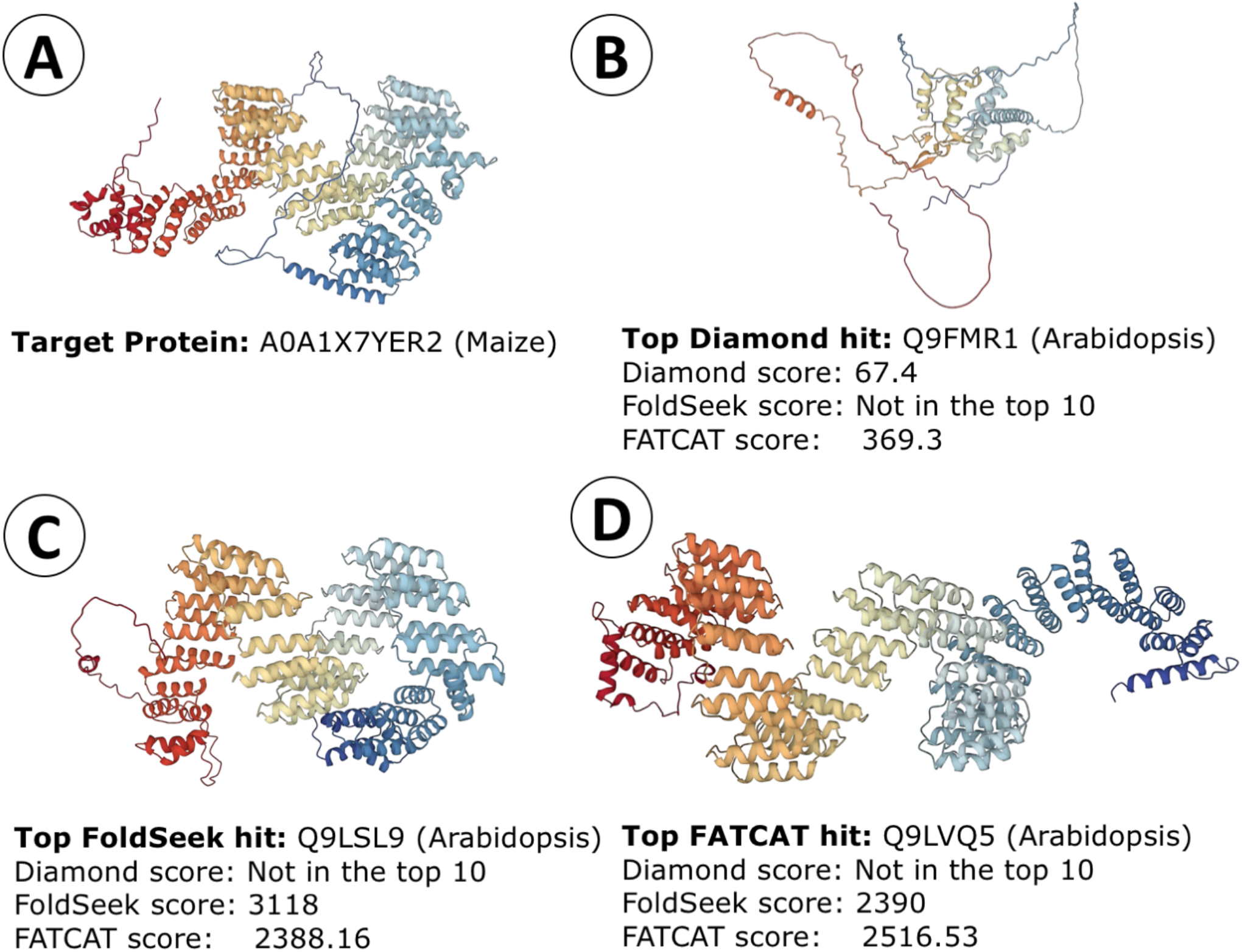
Example of conflicting ortholog predictions. Panel A displays the AlphaFold 3D structure of the maize protein A0A1X7YER2. The three other panels show the AlphaFold structures of the top Arabidopsis hits from Diamond (Panel B), FoldSeek (Panel C), and FATCAT (Panel D). Diamond’s top is Q9FMR1 (AT5G22400) with the functional annotation “Rho GTPase-activating protein 1 (Rho-type GTPase-activating protein 1).” FoldSeek’s top hit is Q9LW84 (At3g16010) with the functional annotation “Pentatricopeptide repeat-containing protein.” FATCAT’s top hit is Q9LSL9 (AT5G65560) with the functional annotation “Pentatricopeptide repeat-containing protein.” Each panel shows the alignment scores of the three methods. If the protein is not in the top ten hits for a method, it is shown as “Not in the top 10.”

An advantage of the FASSO approach is that it assigns confidence values to each predicted ortholog. FASSO uses the results from the reciprocal best approaches to make a final set of orthologs. For example, when using maize as the query proteome, FASSO aggregates the predictions of the individual methods provided in Table 3 to assign confidence labels seen in Table 4. If all three methods predict the same best ortholog pairs, the orthologs are labeled platinum. If two of the three methods agree, the orthologs are labeled gold, and if only a single method makes a prediction, the ortholog pair is labeled silver. As mentioned above, if any method predicts conflicting predictions for a protein, the ortholog pair is moved to a working set. Table 4 shows the count of each confidence level for the final maize predictions and the size of the working sets.

**Table 4:** Maize FASSO results. The table lists the FASSO results when maize was the query proteome. For each of the seven target proteomes, the table lists the count of platinum, gold, and silver orthologs, total FASSO orthologs, and the number of orthologs in the working set. The target proteomes are listed in ascending order based on the time the proteome diverged from a common ancestry with maize.

| Proteome | Platinum | Gold | Silver | Total | Working set |
| --- | --- | --- | --- | --- | --- |
| Sorghum | 10,854 | 3,558 | 1,395 | 15,807 | 8,353 |
| Rice | 8,691 | 3,316 | 1,692 | 13,699 | 7,401 |
| Soybean | 3,108 | 1,673 | 2,415 | 7,196 | 14,255 |
| Arabidopsis | 4,780 | 1,931 | 1,623 | 8,334 | 8,099 |
| Human | 1,058 | 857 | 784 | 2,699 | 2,670 |
| *S. cerevisiae* | 671 | 613 | 355 | 1,639 | 1,865 |
| *S. pombe* | 832 | 680 | 294 | 1,806 | 1,908 |
| Total | 29,994 | 12,628 | 8,558 | 51,180 | 44,551 |

Figure 10 provides stacked bar charts showing the coverage of the maize proteome with annotations from the four approaches for each target proteome. The total length of each bar chart represents the percent coverage of the 39,299 maize proteins with a functional annotation inferred from an annotated ortholog. The color within the bars signifies how FASSO labels these annotations. The bars to the right of the y-axis show platinum, gold, and silver predictions. The bars to the left of the y-axis represent conflicting predictions added to the working set. By design, FASSO will always contain the highest percentage of non-conflicting annotations, which range from over 40% (using sorghum) to below 10% when using the outgroups. Diamond, FoldSeek, and FATCAT may sometimes have higher overall proteome coverage than FASSO, but those predictions often contain lower-quality annotations where the sequence and structure-based methods do not agree. For example, over 50% of the annotations by Diamond, FoldSeek, and FATCAT disagreed with each other when using soybean as the target proteome.

**Figure 10:**
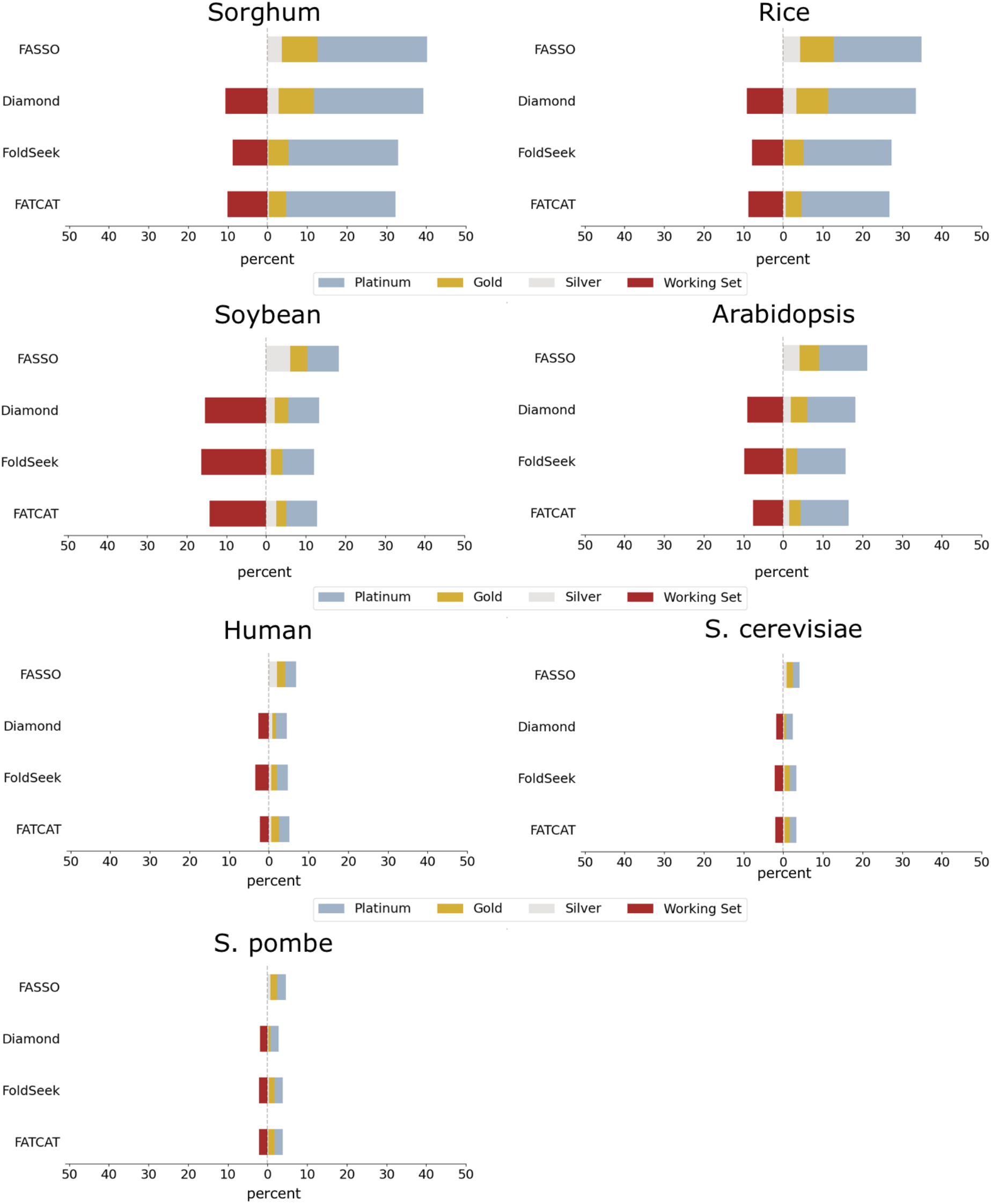
Stacked bar chart of annotated maize proteins. The figure shows seven stacked bar charts showing the percent of proteins (y-axis) in the maize proteome with a functional annotation based on each target proteome. Each chart shows the results of FASSO, Diamond, FoldSeek, and FATCAT. The total length of each bar chart represents the percent coverage of the 39,299 maize proteins with a functional annotation inferred from a predicted ortholog. The color within the bars signifies how FASSO labels these annotations. The bars to the right of the y-axis show platinum, gold, and silver predictions. The bars to the left of the y-axis represent conflicting predictions added to the working set.

As part of the annotation step, FASSO generates Venn diagrams visualizing the frequency of each confidence value in a set of predicted orthologs. Figure 10 compares two example Venn diagrams (maize/sorghum and maize/human). Supplementary Dataset #6 provides the complete set of Venn Diagrams. For the closely related maize/sorghum orthologs, nearly 11,000 platinum orthologs cover over 27% of the maize proteome. Approximately 3,600 gold and 1,400 silver orthologs cover 9.1% and 3.5% of the maize proteome, respectively. For the distantly related pair of maize/human, there are slightly over 1,000 platinum orthologs, less than 3% of the maize proteome. The drop-off of gold and silver level orthologs was less severe at 2.2% for gold and 2.0% for silver. For maize/sorghum, Diamond missed only 2.1% of the 15,807 orthologs, but Diamond did not annotate 34.8% of the 2,700 maize/human orthologs. Table 4 lists the number of orthologs for each confidence level when maize was the query proteome. As the evolutionary distance increased, the number of platinum orthologs decreased quickly (from 10,854 to 671), where the number of gold/silver remained consistent or improved for the plant proteomes.

Additionally, there were more gold/silver annotations for the three outgroups than platinum annotations. The sequence-based ortholog predictions provide the highest coverage for closely related species, but conversely, provide the lowest proteome coverage as the species become more distant. These results illustrate the value of using an aggregate approach, especially for distantly related proteomes.

The catchall term used for annotating proteins with unknown or missing functions is “uncharacterized protein,” or sometimes called “hypothetical protein” [57]. These proteins commonly have evidence of serving a functional role, including having an open reading frame, gene expression, and/or conserved homology. Still, they lack evidence to be assigned a particular function (e.g., functional domains or relationship to orthologs with known function) [58]. These annotations can frustrate researchers looking for valuable clues when searching for causal genes. The percentage of uncharacterized proteins can range from relatively few in a well-annotated proteome to over half the proteome. For example, less than 2% of the human proteins are labeled as uncharacterized while 54% of the sorghum proteome was uncharacterized. Table 2 lists the uncharacterized protein counts for each proteome in this paper. We looked at how well FASSO found new functional annotations to uncharacterized proteins. Table 2 lists the uncharacterized protein counts and the number of uncharacterized proteins with a FASSO annotation. Figure 12 shows a stacked bar chart using this data. The bottom dark green bars represent the percentage of uncharacterized proteins annotated by FASSO (the total number of uncharacterized proteins is listed at the top of the figure). The middle red bars show the percentage of uncharacterized proteins moved to the working set, and the top light green bars indicate the number of remaining unannotated proteins. The human proteome is well annotated; therefore, FASSO could only predict new annotations for 3% of the uncharacterized proteins. For the other proteomes, FASSO predictions had proteome coverages ranging from 18% (*S. cerevisiae*) to 52% (rice). Although the working sets contain conflicting annotations, they also provide a high-coverage set. When we combine the FASSO annotations with the working set, the proteome coverage increases to 22% for *S. cerevisiae* and 72% for rice.

**Figure 11:**
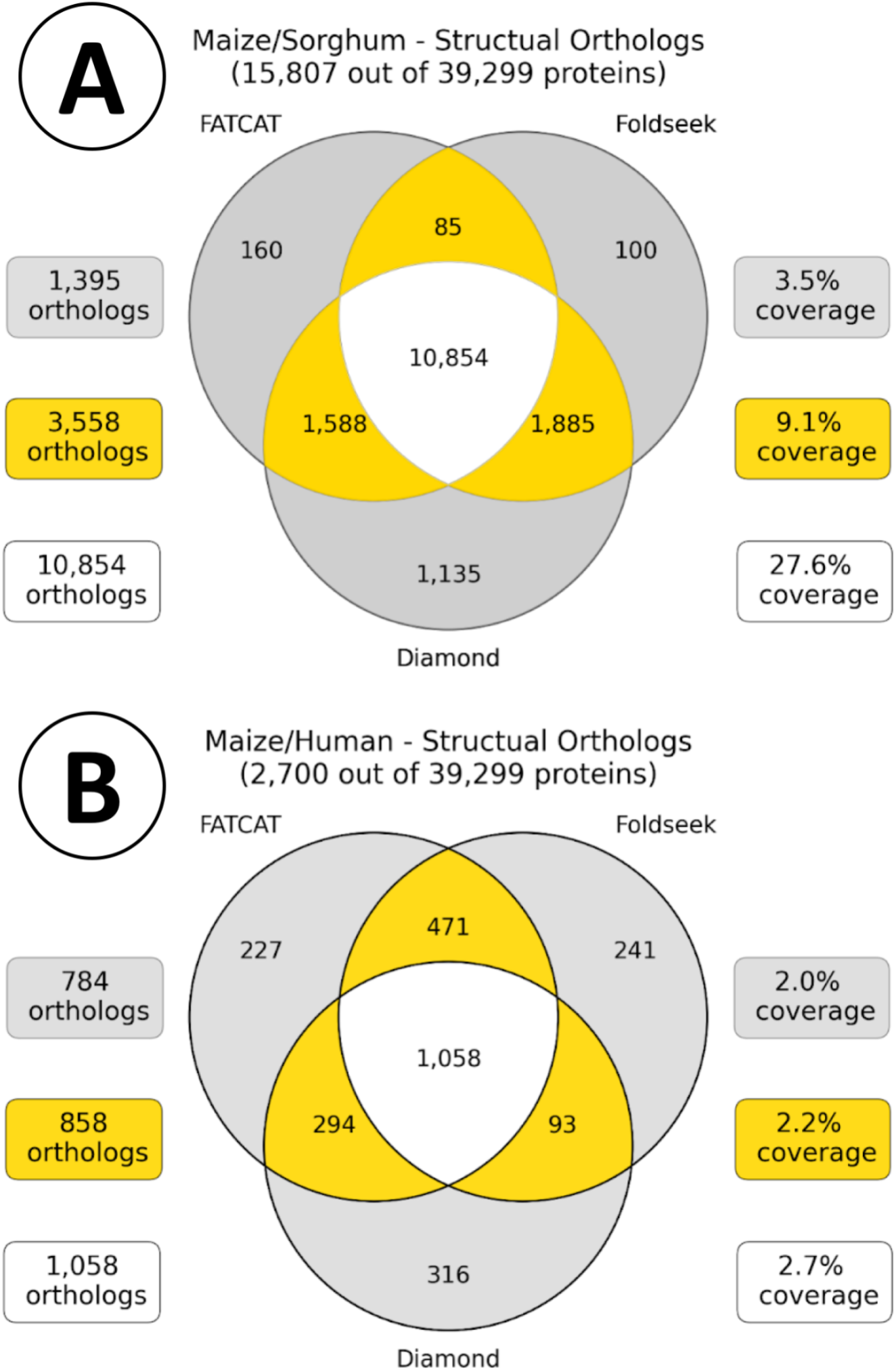
Venn diagram of the FASSO annotation confidence labels for Maize/Sorghum and Maize/Human. FASSO predicts structural orthologs, infers functional annotations based on the predictions, and assigns confidence labels. FASSO generates Venn diagrams to visualize the composition of the confidence labels and how they were generated. Each Venn diagram shows the number of orthologs predicted by Diamond, FoldSeek, and FATCAT and the relationships between those three sets of orthologs. Each type of confidence label count is shown to the left of the diagram and the coverage percentages are shown to the right. Panel A shows the results between the closely related query proteome maize and target proteome sorghum. Panel B shows the results between the distantly related query proteome maize and target proteome human.

**Figure 12:**
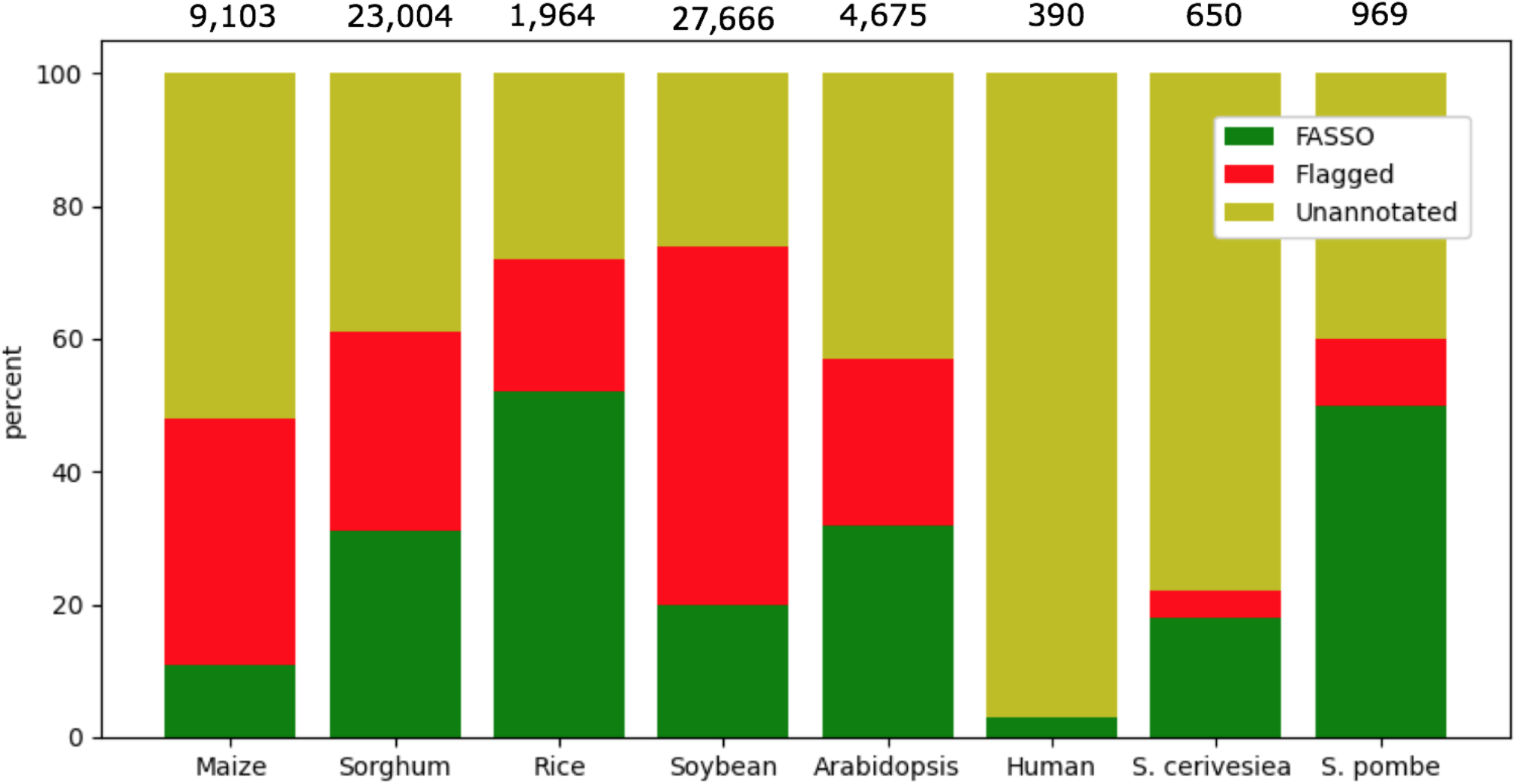
Stacked bar chart of the percent of functional annotations assigned to uncharacterized proteins for each proteome. The x-axis lists each proteome. The total number of uncharacterized proteins for each proteome is listed at the top of the chart. Each bar represents the percentage of those proteins (y-axis) that received an annotation by FASSO (dark green), was flagged and moved to the working set (red), or remained unannotated (light green).

FASSO assigned new functional annotations to 1,021 of the 9,103 uncharacterized proteins in maize. We explored how well gene expression levels support the new functional annotations by reviewing the mRNA abundance levels across 173 samples from 11 studies [42, 59–67] downloaded from qTeller [68] at MaizeGDB [69]. A total of 962 of the 1,021 proteins were assigned a B73 RefGen_v5 [42] gene model based on the MaizeGDB cross-reference assignments [69]. Of these 962 proteins, 90% had expression over 1 FPKM (fragments per kilobase of exon per million mapped fragments) and 72% had an expression of at least 10 FPKM for one or more samples. Supplementary Dataset #7 provides information on the predicted annotations of the uncharacterized maize proteins and the set of FPKM values from qTeller.

Supplementary Dataset #8 contains links to the maize orthologs and annotations at MaizeGDB. Annotating previously uncharacterized maize proteins provides value to researchers looking for functional information about a gene, but additional experimental evidence to validate the annotations is important. Figure 13 highlights three examples of previously uncharacterized maize proteins that are also highly expressed in tissues consistent with their FASSO annotations. In Figure 13A, FASSO annotated the maize protein A0A1D6EGM6 (B73 gene model: Zm00001eb090430) as “Embryonic stem cell-specific 5-hydroxymethylcytosine-binding protein” based on a gold-level annotation in Arabidopsis. 5-Hydroxymethylcytosine (5hmC) is present in specific cell types, including embryonic cells [70], which is consistent with the high levels of expression for Zm00001eb090430 in the tissues “Embryo 16 days after pollination”, “Meiotic Ear” [42], and “Immature unpollinated ear tip”. [66]. Figure 13B displays the expression for protein B4FAY5 (Zm00001eb363710) with a silver-level FASSO annotation of “Anther-specific protein RTS” from rice. RTS (rice tapetum specific) is an anther-specific gene required for male fertility [71]. The gene model Zm00001eb090430 is highly expressed in both the meiotic and immature tassel tissues [42, 66]. A third example (Figure 12C) presents the expression for protein A0A1D6FNG9 (Zm00001eb347250) with a silver-level FASSO annotation of “PMEI domain-containing protein” from soybean. Pectin methylesterase inhibitor (PMEI) genes have shown a role in pollen and anther development in maize [72] and Zm00001eb347250 had the highest expression levels in pollen [73] and anther [62] tissues.

**Figure 13:**
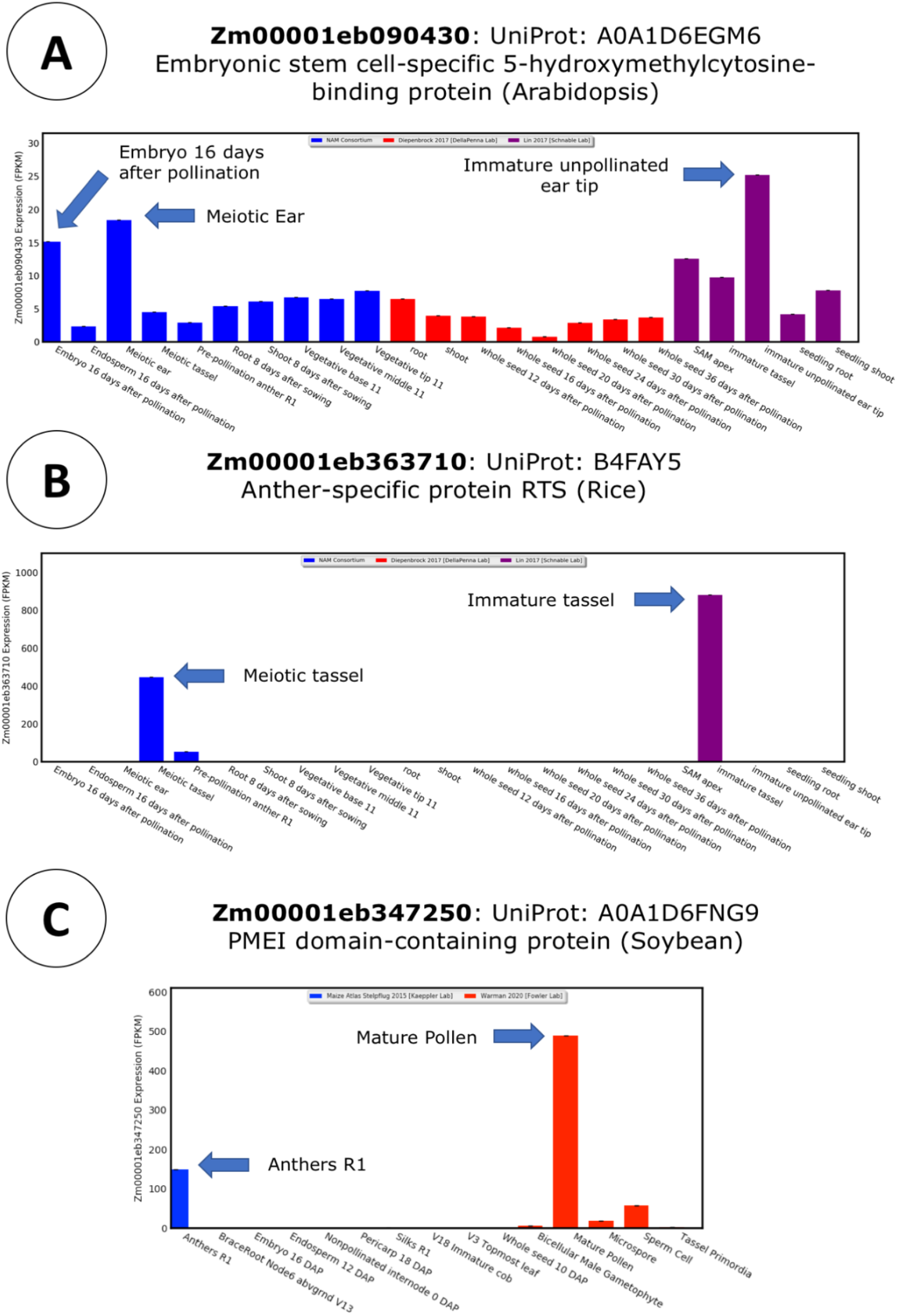
qTeller results for three uncharacterized proteins. The figure shows qTeller gene expression graphs for three maize proteins that have a UniProt annotation of uncharacterized protein, but received a new functional annotation from FASSO. The x-axis lists the gene expression sample name. The y-axis represents the FPKM value for the mRNA abundance expressed by the gene. Blue arrows highlight tissues with a high expression that support the FASSO functional annotation.

These examples exhibit FASSO’s utility in assigning annotations to a broad set of proteins using sequence and structural orthology: A0A1D6EGM6 is annotated based on both sequence (Diamond) and structure (FATCAT) orthology. B4FAY5 is annotated from an ortholog found only in the sequence-based Diamond predictions, and A0A1D6FNG9 is annotated using an ortholog found only with a structure-based method (FATCAT).

## Conclusion

In bioinformatics, sequence similarity has long been used to assign orthologous relationships and infer biological functions. Without protein structure information at a proteome scale, sequence similarity remained an imperfect proxy for functional similarity. Recent advances in protein structure prediction enable us to integrate structure information with previous sequence-based approaches. As sequencing technologies and structural prediction methods improve, new strategies to compare and annotate proteomes will be vital.

This paper introduces FASSO, a tool to functionally annotate proteomes using sequence and structure orthology. FASSO integrates sequence-and structure-based methods to better identify potential orthologs between species. We benchmarked FASSO by predicting structural orthologs and assigning functional annotations for 64 pairs of proteomes across eight species using over 170 billion protein alignments. We compared the sequence-based and structure-based approaches for each proteome pair. Our results exemplify FASSO’s ability to identify additional orthologs, flag orthologs with conflicting predictions, and provide confidence labels to each ortholog. Our method predicted orthologs across a wide range of taxa, including closely related species (diverged less than 15 million years ago) and distantly related species (diverged over 1 billion years ago). We used FASSO to make ortholog predictions to assign over 270,000 functional annotations across the eight proteomes, including new annotations for over 5,600 uncharacterized proteins. Benchmarked using an HPC, our method can produce results within 1-2 days for most proteomes. We present a use-case example of annotating the maize proteome.

The predicted orthologs and functional annotations for maize are available as downloads and genome browser tracks at MaizeGDB.

## Supplementary Data

The supplementary data can be found at the FASSO GitHub repository in the supplementary_data folder: https://github.com/Maize-Genetics-and-Genomics-Database/FASSO/.

Maize data (Dataset #8) is also available at the MaizeGDB downloads page: https://download.maizegdb.org/GeneFunction_and_Expression/FASSO_AlphaFold_Orthologs/.

**Dataset #1 -Diamond, FoldSeek, and FATCAT orthologs and annotations**.

The dataset contains two directories. The “orthologs” directory has the best reciprocal hits from Diamond, FoldSeek, and FATCAT for every pairwise combination of proteomes. The “annotations” directory contains a merged tab-separated file showing the UniProt annotations from the predicted orthologs from Diamond, FoldSeek, and FATCAT. The data is in the GitHub repository under /supplementary_data/Dataset_1_diamond_foldseek_fatcat_annotations/.

**Dataset #2 -FASSO structural orthologs**.

The dataset contains the FASSO orthologs, working sets, and Venn diagram data file for every pairwise combination of proteomes. The data is in the GitHub repository under /supplementary_data/Dataset_2_FASSO_orthologs/.

**Dataset #3-FASSO functional annotations**.

The dataset contains the FASSO UniProt annotations assigned to the proteins in each proteome. The data is in the GitHub repository under /supplementary_data/Dataset_3_FASSO_annotations/.

**Dataset #4-Diamond, FoldSeek, and FATCAT top 10 ortholog predictions**.

The dataset contains the top 10 hits from Diamond, FoldSeek, and FATCAT for every pairwise combination of proteomes. The data is in the GitHub repository under /supplementary_data/Dataset_4_top_ten_hits/.

**Dataset #5-Diamond, FoldSeek, and FATCAT heat maps**.

The dataset contains the heat maps that compare the top 10 ranked hits between Diamond/FATCAT, FoldSeek/FATCAT, and Diamond/FoldSeek. The data is in the GitHub repository under /supplementary_data/Dataset_5_heat_maps/.

**Dataset #6-Venn diagrams of the confidence labels of the FASSO ortholog predictions**. The dataset contains Venn diagram images showing the composition of the confidence labels for the FASSO ortholog predictions for each proteome. The data is in the GitHub repository under /supplementary_data/Dataset_6_venn_diagrams/.

**Dataset #7-FASSO annotations of the uncharacterized maize proteins**.

The dataset contains the FASSO annotations for the maize proteins with “uncharacterized protein” annotations in UniProt. The data is in the GitHub repository under /supplementary_data/Dataset_7_FASSO_uncharacterized_annotations/.

**Dataset #8-Maize orthologs and functional annotations available at MaizeGDB**.

The dataset contains all the maize datasets from this paper including FASSO orthologs, FASSO annotations, Venn diagrams, FASSO annotations for uncharacterized proteins, Diamond, FoldSeek, and FASTCAT top hits, orthologs, and annotations. An additional file shows FASSO annotations for the maize uncharacterized proteins with gene expression levels for each protein. The data is in the GitHub repository under /supplementary_data/Dataset_8_maize_annotations/.

## Acknowledgments

The work for this project was performed on the Ceres high-performance cluster as part of the USDA-ARS SCINet initiative. We would like to thank the SCINet administrative staff and the Virtual Research Support Core team.

## Author Contributions

CMA and MRW conceived, designed, and directed the study. CMA was responsible for software design, validation, visualization, and data analysis. RKA, LCH, and JMG provided data curation, guidance on the maize use-case study, and testing. JLP and EKC gave technical guidance, reviewed the code, and integrated resources at MaizeGDB. CMA, TZS, and MRW led funding acquisition and project administration. CMA and SS drafted the manuscript. The final version of the manuscript was generated with input and contributions from CMA, SS, RKA, JLP, EKC, LCH, JMG, TZS, and MRW. All authors approved the final version of the manuscript.

## Data Availability Statement

Publicly available datasets were analyzed in this study. This data can be found here at the EBI AlphaFold database and the UniProt Knowledgebase (UniProtKB) for the following species: *Arabidopsis thaliana* (UP000006548), *Glycine max* (UP000008827), *Homo sapiens* (UP000005640), *Oryza sativa* (UP000059680), *Saccharomyces cerevisiae* (UP000002311), *Schizosaccharomyces pombe* (UP000002485), and *Zea mays* (UP000007305). The datasets for *Sorghum bicolor* were downloaded from the Google Cloud with taxonomy id: 4558. The code developed for this project and supplementary data can be found on GitHub https://github.com/Maize-Genetics-and-Genomics-Database/FASSO.

